# How to catch a blank mind? Brain similarities and differences between self- and probe-caught mind blanking

**DOI:** 10.64898/2026.06.10.731344

**Authors:** Agustina Aragón-Daud, Paradeisios Alexandros Boulakis, Sepehr Mortaheb, Gilles Vandewalle, Fabienne Collette, Kaustubh R. Patil, Athena Demertzi, Federico Raimondo

## Abstract

Mind Blanking (MB) is a mental state characterized by the experience of a seemingly empty mind. This apparent mental emptiness challenges the theoretical view of a continuous and content-full stream of thought, leading to the question of how it is possible to capture MB events in the first place. MB research has relied largely on probe-caught experience-sampling, during which participants are interrupted at random times to report their mental state. This approach risks missing some MB events as they may occur between probes. Self-caught paradigms, on the other hand, allow participants to report MB upon spontaneously realizing it, relying on meta-awareness. What remains unclear is whether these two methods capture similar properties of MB. To address this gap, we investigated whether probe-caught MB (pMB) and self-caught MB (sMB) methodologies converge on behavioral and brain correlates. Twenty-two participants underwent 3T fMRI scanning while performing both a pMB and a sMB experience-sampling task. We found that both MB reports were uniformly distributed over time and that their report frequencies significantly correlated across individuals, pointing towards a behavioral convergence across the two approaches. Time-varying functional connectivity further revealed that both pMB and sMB were linked to a hyperconnectivity pattern, and that sMB had higher global signal magnitude compared to mind wandering, pointing towards a brain mode characterized by low arousal levels. According to our predictions, sMB elicited greater activations in the anterior cingulate cortex and (pre)motor areas, as well as deactivations in occipital, temporal, and frontal regions, compared to pMB, potentially reflecting metacognitive processes involved in self-detecting MB, alongside motor preparation associated with self-initiated responses. Together, our results show an overall converge of sMB and pMB on neurobehavioral correlates. This positions the self-caught methodology as a complementary tool for studying MB without external interruption, while highlighting the role of meta-awareness in self-report detection.

## 1. Introduction

While wakefulness is typically characterized by a continuous stream of thought, individuals can experience periods of a seemingly empty mind. This experience is linked to a mental state known as Mind Blanking (MB), formally defined as a waking state in which individuals report no mental content or are unable to describe their immediate experience (Andrillon et al., 2025; Boulakis & Demertzi, 2025; Ward & Wegner, 2013). So far, research into MB has primarily relied on probe-caught methodologies, where participants are interrupted to report their immediate mental state at random moments (e.g., “What was on your mind just now?”). However, individuals can also spontaneously catch themselves in a blank state without an external prompt (i.e., self-caught). To the best of our knowledge, only one study so far has used the self-caught paradigm to characterize MB behaviorally (Ward & Wegner, 2013), while their brain correlates have not yet been explored. Consequently, it remains unknown whether these two reporting methodologies capture the same neurobiological phenomenon.

Probe-caught methods have been extensively used both during resting state (Boulakis, Simos, et al., 2025; Koroma et al., 2023; Unsworth et al., 2021; Van Calster et al., 2017) and during cognitive tasks (Andrillon et al., 2021; Huijser et al., 2020; Robison et al., 2019, 2020; Unsworth et al., 2025). These studies have shown that probe-caught MB (pMB) events are reported sparsely (Boulakis et al., 2026), but uniformly across probes (Mortaheb et al., 2022). Converging evidence from neuroimaging studies points towards reduced cortical excitability and hyper-synchrony during pMB. First, the brain elicits parietal slow wave activity during pMB reports that, despite its localized nature, morphologically resembles the slow waves observed globally during deep sleep (Andrillon et al., 2021). Second, the brain shows widespread deactivations across cortical areas (Boulakis et al., 2023). Third, functional connectivity (FC) during pMB self-organizes in a hyperconnected pattern where all regions are synchronized together (Mortaheb et al., 2022) as also observed in unconscious states (Aedo-Jury et al., 2020; El-Baba et al., 2019). However, this relationship seems to be mediated by perceived arousal and task context. For instance, during a sustained attentional task, pMB has been associated with the typical anticorrelated FC pattern observed in wakefulness across vigilance levels, yet was associated with the hyperconnected pattern only during self-reported high vigilance levels, potentially due to hyperconnectivity being more dominant across mental states during sleepiness (Boulakis, Kusztor, et al., 2025). Lastly, pMB reports increase in states of low arousal, such as after sleep deprivation (Boulakis, Simos, et al., 2025), and, compared to other mental states, are characterized by higher fMRI global signal (GS) amplitude (Mortaheb et al., 2022), which widely reflects low cortical arousal levels (Liu et al., 2017; Wong et al., 2013).

Even though probe-caught methods are valuable, they possess an inherent limitation: they rely on externally timed cues and may miss episodes that occur between probes. Indeed, MB in probe-caught methodologies is typically captured by 10% of the probes (Boulakis et al., 2026), rendering it a highly time-consuming methodology. Approaches that do not depend on external sampling could, therefore, offer a more efficient alternative by capturing the entire sampling period and allowing for the unconstrained accumulation of reports. A promising alternative is the self-caught paradigm, which has been widely applied in mind wandering (MW) research (Chu et al., 2023) but rarely to MB (Ward & Wegner, 2013). In this method, participants are instructed to report whenever they notice or ‘catch’ themselves in a specific mental state. By design, this approach captures MB events with meta-awareness (Chu et al., 2023), meaning one must realize their mind feels empty in order to report it (i.e., awareness of absence) (Mazor & Fleming, 2020). Given that this introduces a distinct metacognitive demand, and because the paradigm has rarely been applied to MB, it remains unknown whether the self-caught MB (sMB) shares behavioral and brain correlates as pMB, or if the act of self-catching fundamentally alters its phenomenological and neurophysiological correlates.

The present study aims to compare self-caught and probe-caught methodologies and to investigate whether sMB shares the established behavioral properties and neural correlates of pMB. While we expect the core signature of this state to be similar, we hypothesize that sMB recruits additional brain regions related to meta-awareness and metacognitive monitoring.

## 2. Methods

### 2.1. Participants

Twenty-two individuals participated in the study (45% female; mean age = 29 ± 4 years). Participants were recruited via campus poster advertisements and intranet electronic invitations. All participants provided written informed consent in accordance with the Declaration of Helsinki. The study was approved by the Ethics Committee of the University Hospital of Liège (Nr: B707201941887). Exclusion criteria involved MRI contraindications and history of neurological disorders. All participants were right-handed and french speakers with academic backgrounds.

### 2.2. Design & Tasks

The study employed a within-subject design, in which participants performed two experience-sampling tasks in a 3T MRI scanner (Magnetom Prisma, Siemens Medical Solutions, Erlangen, Germany): one with the self-caught, and the other with the probe-caught methodology. Tasks were performed in two sessions separated by a 15-minute break. The first session included the self-caught task, alongside other cognitive tasks outside the scope of the present study. The second session comprised the probe-caught task and diffusion-weighted imaging, which is also outside the scope of this study. In the self-caught task (approx. 8 min duration), participants were asked to press a button whenever they felt they had an MB event. In the probe-caught task (approx. 19 min duration), participants were interrupted at random times by auditory and visual probes (n=25, inter-probe interval: 30-60s), inviting them to use a button press to report their immediate mental state. Participants were asked to report if their thoughts were in the “past”, “future”, “present”, or simply “nowhere” (MB). When participants choose any of the first two options, they were presented with a second question asking if they were thinking of themselves or someone else (all coded as MW). When they selected “present” they were asked if their thought was stimulus dependent (Sensations) or independent (MW). This multi-step questioning tree was designed to mirror the specific dimensions of time, self-referentiality, and stimulus-dependence manipulated across the broader experimental protocol. However, because these granular distinctions are outside the scope of the present study, reports were collapsed into broader categories (MB, MW, and Sensations) for analysis. Frequency of other mental states aside from MB can be found in S1. If a response took longer than 10 seconds, it was coded as an error and excluded from the analysis.

The tasks were implemented in MATLAB using the Psychophysics Toolbox (PTB) (Brainard, 1997) and are available at https://gitlab.uliege.be/poc/self-caught-mb. Experimental design and analysis are illustrated in Fig 1. During both tasks, participants were in a resting state, instructed to let their mind wander, with eyes open, fixating on a black screen. For each task, we defined rest and mental state periods. For the probe-caught task, mental states pMB, MW, and Sensations) were defined as the 10s before the probe (Boulakis, Kusztor, et al., 2025; Mortaheb et al., 2022; Van Calster et al., 2017). For the self-caught task, sMB was defined as the 10s before the button press.

**Fig 1.**
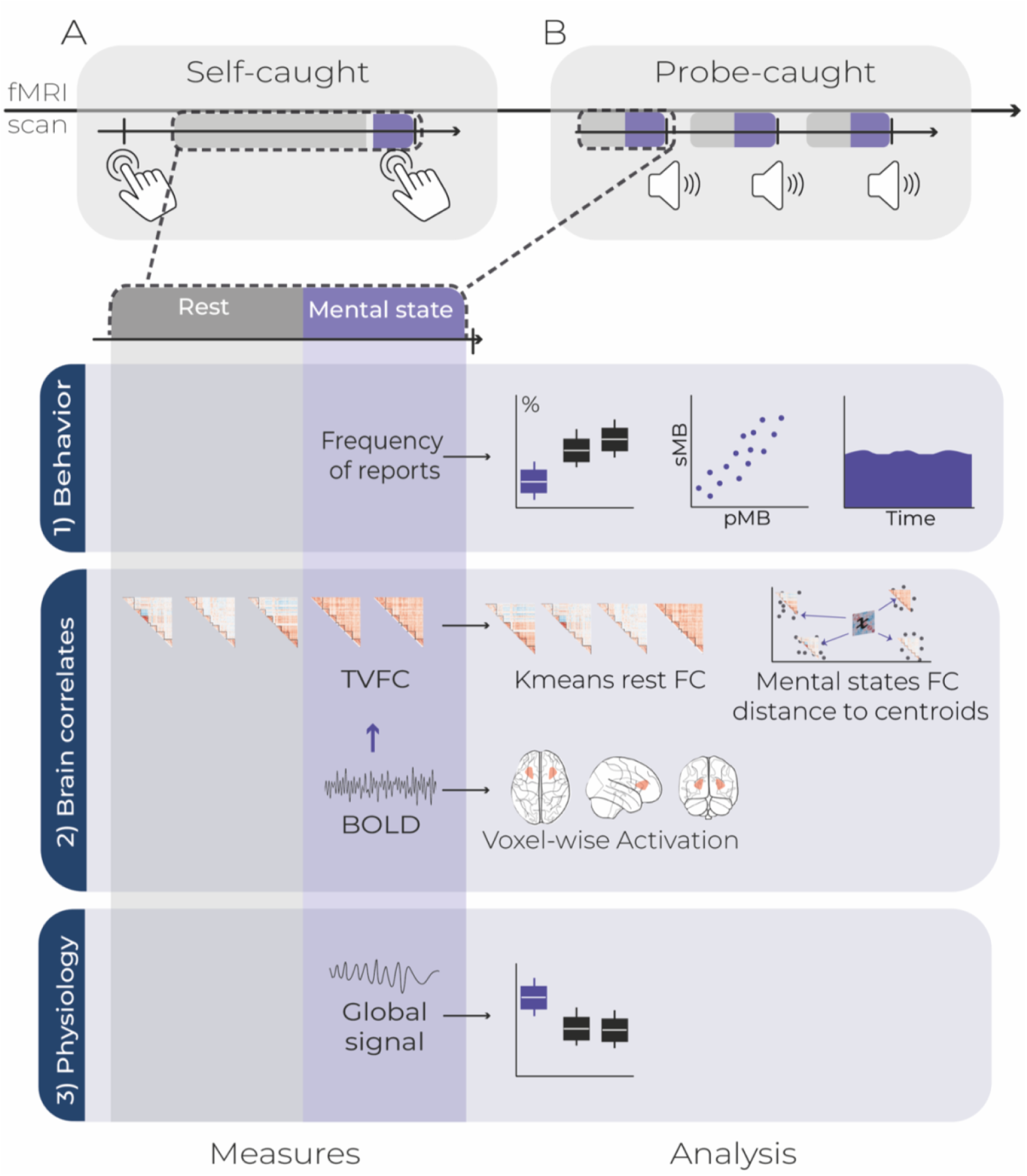
Experimental setup and analysis. Twenty-two participants performed A) self-caught and B) probe-caught experience-sampling tasks while lying down in a 3T MRI scanner. In both tasks, mental states epochs (in blue) were defined as the 10 s preceding the self-report button press (self-caught) or auditory and visual probe (probe-caught), and rest epochs as 30 s (in grey) of uninterrupted resting activity (which could be longer in the self-caught task if no button press occurred). Analysis spanned three domains: (1) Behavior: We analyzed report frequencies, cross-task correlations across individuals, and the temporal distribution of MB reports. (2) Brain correlates: We extracted BOLD timeseries for two distinct analyses. First, we computed time-varying functional connectivity (TVFC) across both tasks; we clustered rest timepoints to identify representative connectomes (centroids) and calculated the distance of the TVFC patterns during each mental state deviated to these baseline rest clusters. Secondly, a voxel-wise activation analysis contrasted sMB versus pMB. (3) Physiology: We tested whether mental states differed in global signal (GS) magnitude. TVFC = time-varying functional connectivity; BOLD = blood oxygen level dependent; GS = global signal.

Regarding rest periods, these were defined as up to 30s in between probes in the probe-caught task. In the self-caught task, rest timepoints were defined as periods of uninterrupted resting state (i.e., no button press), with no specific length. To avoid signal contamination, timepoints surrounding each sMB report were excluded (buffer of 40s sMB report: 30s before and 10s following the button press).

### 2.3. fMRI

#### 2.3.1. Functional Imaging parameters

FMRI data were acquired using a standard transmit–receive quadrature head coil and included T2*-weighted gradient-echo echo-planar imaging (EPI) sequence using the following parameters: repetition time (TR) = 1170 ms, echo time = 30 ms, field of view = 216 × 216 mm², acquisition matrix = 72 × 72 (3 × 3 mm² in-plane resolution), and 36 axial slices of 3 mm thickness with a 25% interslice gap (0.75 mm), covering most of the brain. Slice acquisition was performed in an interleaved order, with phase encoding in the anterior-to-posterior direction. Multi-band acceleration factor of 2 was applied to reduce acquisition time. A total of 1000 volumes were acquired per run. A high-resolution T1-weighted magnetization prepared rapid gradient echo image (MPRAGE) was acquired for anatomical reference (TR = 1960 ms, echo time = 2.19 ms, inversion time = 900 ms, field of view = 256 × 240 mm^2^, matrix size = 256 × 240 × 176, voxel size = 1 × 1× 1 mm^3^). Participants’ heads were comfortably secured with a vacuum cushion to reduce movement during scanning. Visual stimuli were presented on a screen located at the back of the scanner, seen via a mirror attached to the head coil.

#### 2.3.2. fMRI Preprocessing

FMRI data were preprocessed using fMRIPrep (version 24.1.1; (Esteban et al., 2018)), a standardized pipeline for robust fMRI preprocessing: for each participant, functional data were corrected for susceptibility distortion using dual-echo gradient-recalled echo (GRE) field maps, aligned to their native T1-weighted (T1w) anatomical space and spatially normalized to the MNI152NLin2009cAsym template at 2 mm isotropic resolution. Anatomical preprocessing, including brain extraction and normalization, was performed using ANTs (2.5.0; (Avants et al., 2014)). Tissue segmentation of cerebrospinal fluid, white matter, and gray matter was performed using FSL (version 6.0.4; (Jenkinson et al., 2012)). As surface-based analyses were not required (parcellation was conducted in volumetric space using the predefined Schaefer atlas), full cortical surface reconstruction was skipped. Given the short TR (<2 seconds) and to avoid altering the temporal autocorrelation structure important for estimating time-varying FC, slice timing correction was not applied.

### 2.4. Analysis

#### 2.4.1. Behavioral properties of MB

To assess the consistency of individual MB reports across conditions, we computed the Spearman correlation between the number of MB reports per subject for each task. The Kendall Rank coefficient was also computed for complementary robustness. This analysis aimed to evaluate whether MB occurrence reflects a stable individual characteristic, whereby participants who report more MB in one task also tend to report more in the other. Following previous studies (Mortaheb et al., 2022) we also examined the temporal distribution of MB reports within each task. Acquisition time was divided into 5 equal-duration bins, and the number of MB reports in each bin was counted. We then compared the resulting distribution against a uniform distribution using a χ² test, and calculated effect size using φ (φ = √(χ²/n)). Additionally, to test for evidence in favor of the null hypothesis (uniformity), we computed a Bayes factor comparing a uniform distribution (all bins equally likely) against a non-uniform distribution multinomial model with a symmetric Dirichlet(1) prior, where BF₁₀ < 0.33 indicates evidence for uniformity (null hypothesis), whereas values between 0.33 and 1 were considered inconclusive.

#### 2.4.2. Coupling between functional connectivity patterns and mental states

To estimate functional connectivity patterns, first preprocessed BOLD signals were denoised using junifer (version 0.0.7.dev47; (Mandal et al., 2023)). Confound regression included 24 motion parameters, white matter and CSF signals. Time series were detrended, z-scored, and bandpass filtered (0.008-0.09Hz). A gray matter mask was applied using a probabilistic threshold of 20%. The brain was parcellated according to the Schaefer-100 atlas (100 regions, 7-network version) (Schaefer et al., 2018), and instantaneous phase connectivity (IPC) was computed at each time point using a second-order Hilbert transform after re-filtering the data between 0.01 and 0.04 Hz. To normalize the distribution for statistical analysis, IPC values were Fisher z transformed.

Following previous studies (Boulakis, Kusztor, et al., 2025; Mortaheb et al., 2022), we identified recurring connectivity patterns by applying k-means clustering to resting state IPC matrices using cosine distance (*n* rest IPC = 17.687 matrices; 60% from probe-caught and 40% from self-caught task; additional analysis to guarantee same resting state baselines can be found in S2). While internal validation metrics did not converge on a single optimal *k* (S3), we selected *k*=4 based on established literature identifying four stable resting-state pattens (Demertzi et al., 2019; Mortaheb et al., 2022). We confirmed the stability of our findings by repeating the analysis for k=3 through k=6 (see S4). Resulting centroids were ordered by their spatial standard deviation (a proxy for pattern metastability) (S1). We estimated the occurrence rate of each pattern counting the number of matrices assigned to each centroid for each participant. We then compared occurrence rates with a paired t-test with *p*-FDR < 0.05 corrected for multiple comparisons (S5).

To determine if the connectivity pattern of a mental state (sMB, pMB, MW and Sensations) was associated with a specific baseline network configuration, we calculated the cosine distance at the single-timepoint level. Specifically, for a single sample (timepoint) of each report, we computed the cosine distance between that mental state’s IPC matrix and each of the four canonical connectivity centroids derived from the resting state timepoints. Distances to each cluster were modeled using a Bayesian Generalized Linear Mixed Model. To account for the positive-bounded and negatively skewed nature of the distance metrics, we modeled the response variable using a Skew-normal likelihood. Population-level parameters were assigned weakly informative Skew-normal priors (*ξ*=0, ω =1, α = -2), while group-level standard deviations were modeled using a regularizing Half-Cauchy prior (scale = 0.5). The model included mental state reports as fixed effects and subject-specific random intercepts. Model fit was compared against a null model (containing only intercept and random effects) using Bayes Factors, Akaike Information Criterion (AIC) and Bayesian Information Criterion (BIC) (S6). Bayes Factor favored the null model over the mental state model for each cluster, likely due to the inherent inter-subject variability of spontaneous cognition and the parsimony-based penalties applied by Bayes Factors. Nonetheless, AIC and BIC supported the response model. To explore even marginal effects of the mental states, we performed planned post-hoc comparisons using Estimated Marginal Means between the distances of each mental state to each pattern (S7). A contrast was considered supported if its 89% credible interval excluded zero (Makowski et al., 2019).

#### 2.4.3. Global signal (GS) amplitude as a proxy to arousal estimation across mental states

We considered GS as the average BOLD activity across whole-brain voxels (Fox et al., 2009). As mentioned, GS amplitude serves as an indirect proxy for arousal levels, where higher amplitude typically indicates lower arousal and vice versa (Liu et al., 2017; Wong et al., 2013). Given that MB is often associated with low arousal, we investigated whether GS magnitude (i.e., the L2 norm of the global signal time course during each mental state) differed between mental state reports.

The GS time series were extracted for each subject and task using the standard fMRIPrep pipeline. To account for potential baseline shifts between task sessions (self-caught vs probe-caught task), we performed a session-specific baseline normalization. For each task, the raw GS was divided by the mean GS during that session’s rest periods. To account of the sensibility of the L2 norm (signal magnitude) to the number of data points, all events were truncated to a uniform distribution of 9 timepoints. We calculated the L2 norm for each event to obtain a single GS magnitude value per report.

GS magnitude was modeled using a Generalized Linear Mixed-Effects Model, including mental state reports (sMB, pMB, Sensations, MW) as a fixed effect and a random intercept per subject. Given that GS magnitude values are strictly positive, we utilized a gamma distribution with a log link function. Post-hoc comparisons between mental states were performed using marginal means. All reported p-values were adjusted for multiple comparisons using the Tukey method. We additionally tested whether the global-signal effect could be explained by task duration by including total timepoints as a covariate (S8).

#### 2.4.4. Voxel-wise activation differences between MB report types

We employed a mass-univariate approach based on a General Linear Model (GLM) to identify voxels exhibiting significant activation differences between MB types. Given that only 13 participants reported MB in both tasks, and by considering the limited number of reports per participant, we adopted a fixed-effects approach to maximize statistical power, by pooling all available events across the sample. As a consequence, the resulting statistical inference is valid for this specific sample, rather than generalizable to the broader population. All subjects’ BOLD data was entered into a single first-level GLM. Because the probe-caught and self-caught events were recorded in separate sessions, each subject contributed two runs. The design matrix included the mental state reports (sMB, pMB, Sensations, MW), task regressors (probe or button press) and rest periods, convolved with a canonical SPM hemodynamic response function. The nuisance regressors included six motion parameters and a binary motion outlier regressor for volumes exceeding framewise displacement of 0.5 mm. A cosine-basis drift model was used to implement high-pass filtering (>0.008 Hz). Signal intensity was grand mean scaled to ensure that beta estimates reflect the percentage of signal change. To account for temporal autocorrelation in the BOLD signal, we employed a first-order autoregressive model. Data were spatially smoothed using a Gaussian kernel of 6 mm FWHM. Then, we computed the contrast [sMB > pMB] as the primary contrast of interest. To control for multiple comparisons, we applied voxel-wise false discovery rate (FDR) correction at *p*<.05 to the resulting z-statistic map. For transparency purposes (Taylor et al., 2023), results were visualized on a brain template using transparent thresholding, with the statistical map visualized at an uncorrected threshold of *p*<0.001 (z = 3.29). FDR-corrected significant voxels were outlined with black contours.

## 1. Results

### 1.1. MB reports converged on their behavioral properties across the two methods

To assess the reliability of MB across self-report methods, we compared the frequency of episodes captured via self-caught and probe-caught tasks. Participants reported on average 5 sMB (range = 0-31, N = 119) and 2 pMB episodes (range = 0-7, N = 55). In terms of method-specific relative frequency, pMB episodes were reported in 10.8% of total probes, while sMB reports occurred at a rate of approximately 1 episode per 2.54 minutes of acquisition. A significant positive correlation was observed between sMB and pMB frequencies (r = .49, *p*=.019; τ = .39, *p*=.019, Fig 2A), suggesting MB reporting was consistent at the individual level.

**Fig 2.**
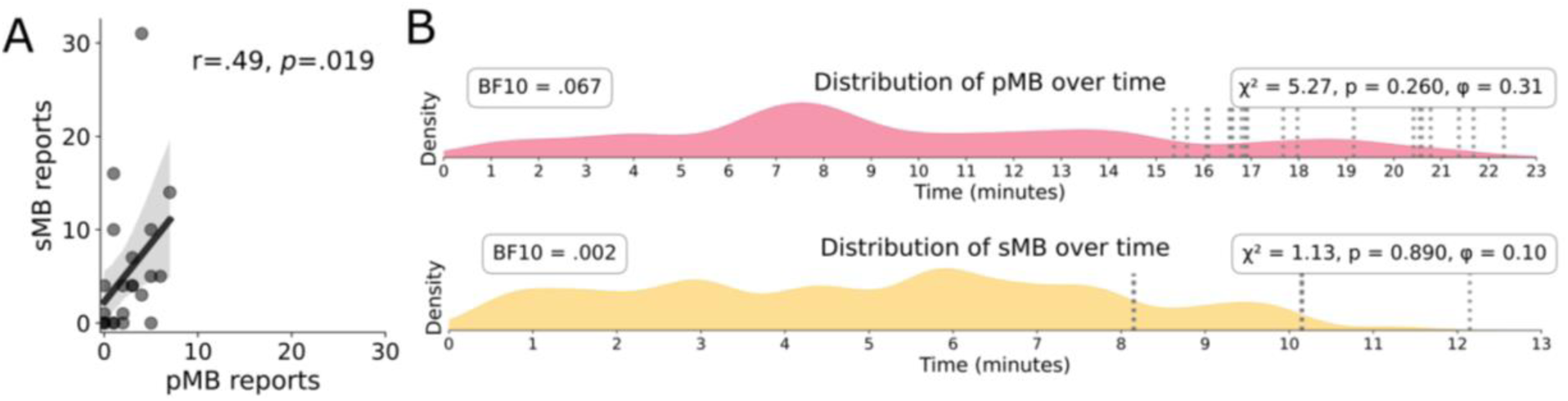
Consistent rate and uniform distribution of MB reports across tasks and time. A) The positive Spearman (r) and Kendall (τ) correlation between the number of sMB and pMB reports per subject indicate that the two report methods are reflecting similar content at the individual level. B) The temporal distribution of pMB (upper panel) and sMB (lower panel) reports during each task indicated that the MB reports were uniformly distributed across both tasks as evidenced by the Bayes factor supporting the null hypothesis. Vertical dashed lines indicate each subject’s task end time. sMB = self-caught mind blanking; pMB = probe-caught mind blanking.

The temporal distribution of MB reports was analyzed to determine if the occurrence of these episodes was influenced by time (Fig 2B). The distribution of MB reports across acquisition time did not significantly deviate from a uniform distribution in either task. Furthermore, Bayes factor analyses yielded evidence supporting the null hypothesis, supporting that the MB reports were uniformly distributed across both tasks.

### 1.2. MB reports share a connectome signature of global functional hyperconnectivity

To identify the primary brain states that emerge during rest, we applied k-means (k=4) clustering to the time-varying FC of the rest timepoints. Fig 3A show the four identified distinct FC patterns. Both Pattern 1 and Pattern 2 showed anticorrelation between the DMN and bottom-up networks, including attentional and sensory networks. Pattern 1 additionally showed anticorrelation between bottom-up and control network (CN), together with high within-somatosensory network connectivity. Pattern 3 showed a generalized low connectivity, characterized by reduced connectivity both within and between networks. Finally, Pattern 4 displayed an all-to-all positive connectivity (hyperconnectivity) within and between networks. The generalized low connectivity (Pattern 3) occurred significantly more often than all the other brain patterns (Pattern 1 – Pattern 3: t = -6.88, FDR-*p <* .0001; Pattern 2 – Pattern 3: t =-6.09, FDR-*p* < .0001; Pattern 3 – Pattern 4: t = 4.12, FDR-*p* = .001; all other comparisons FDR-p < .05, Fig 3A).

**Fig 3.**
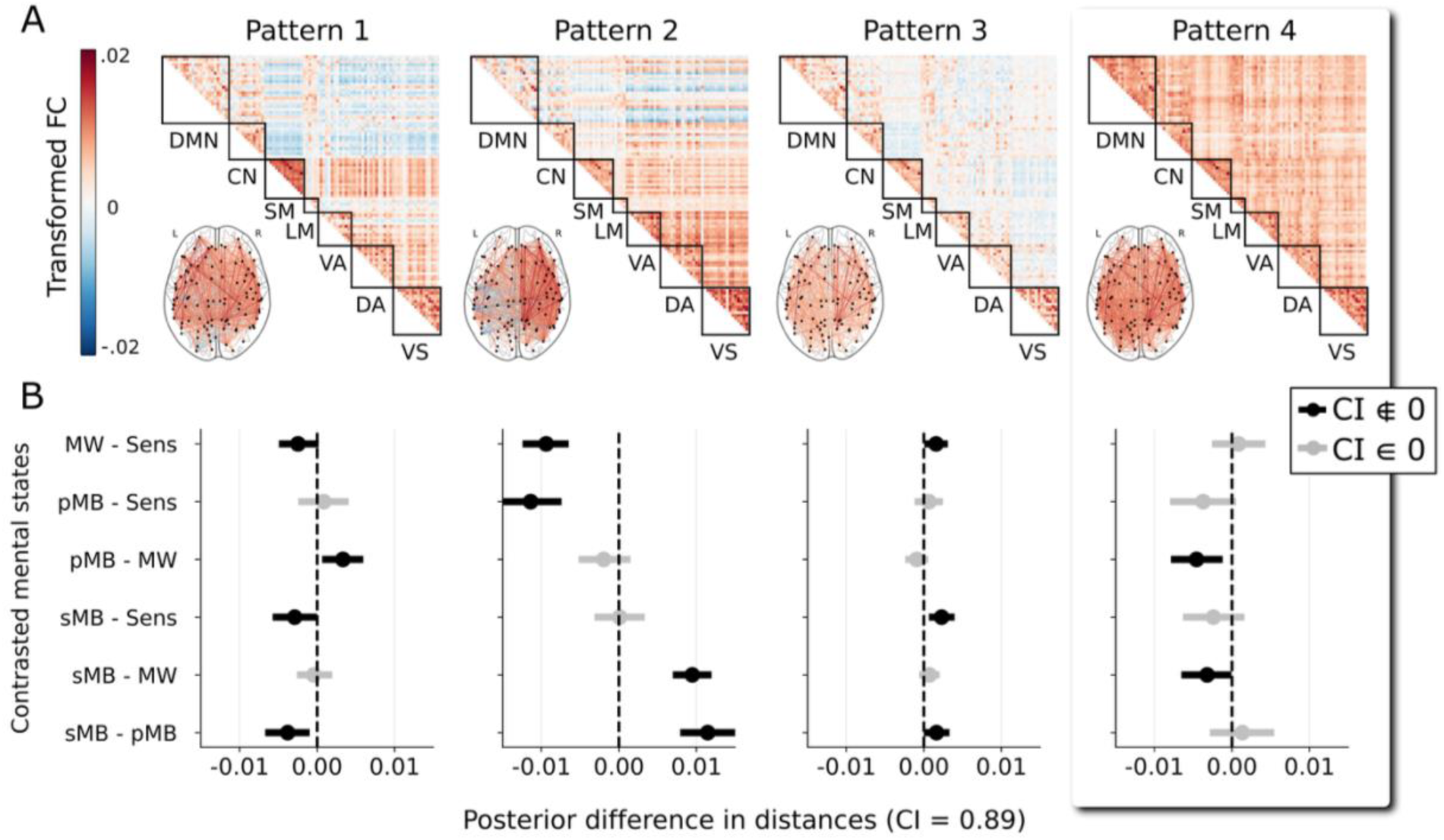
Mind Blanking reports show higher similarity to a hyperconnected pattern of functional connectivity both during the self-caught (sMB) and probe-caught (pMB) method. A) Data-driven k-means centroids of time-varying functional connectivity and connectome indicates that the brain self-organizes into a pattern of anticorrelations between the DMN and sensory networks (Pattern 1), a pattern of somatosensory integration (Pattern 2), a pattern of lower overall coherence pattern (Pattern 3) and a pattern of hyperconnectivity (Pattern 4). Scale shows Fisher Z and L2 normalized functional connectivity. Positive values indicate higher phase synchronization, while values close to zero indicate the absence of a systematic phase coherence. B) Estimated Marginal Means contrast between distances of each mental state functional connectivity matrices to each centroid. Contrast estimate on the right side means the first mental state of the contrast is closer to that pattern. When comparing contrasts across all patterns, both MB reports systematically had higher similarity (lower distance values) to the hyperconnected connectome (Pattern 4) compared to mind-wandering. Significant contrasts are black-colored. DMN = default mode network; CN = control network, SM = somatosensory network, LM = limbic network, VA = ventral attentional network, DA = dorsal attentional network, VS = visual network, MW = mind wandering, Sens = sensations, pMB = probe-caught mimd blanking, sMB = self-caught mind blanking.

Having defined the FC self-organization during rest, we next tested whether the reported mental states showed greater proximity to any of these specific configurations. Overall, the fit indices (BF, BIC, AIC) did not converge on a single preferred model across all patterns (S6). Given that Pattern 4 has been previously associated with pMB during rest (Mortaheb et al., 2022), we focused our primary analysis on this hyperconnected state. Consistent with our hypothesis, sMB and pMB time-varying FC matrices showed significantly higher similarity to this hyperconnected state than mind wandering (sMB-MW: Δ = -0.003, 89% CI [-0.006 -0.001], p < .0001; pMB-MW: Δ = - 0.004, 89% CI [-0.007 -0.002], p < .0001) but did not differ significantly from sensations (*p* > .05). Overall, these findings suggest that both forms of MB episodes share a common FC signature characterized by global hyperconnectivity, regardless of the self-report detection method.

Further analysis revealed additional differences in the proximity of other mental states to other patterns (Fig 3C, S7). Patterns 1 and 2 showed similar proximity profiles: for Pattern 1, MW and sMB were significantly closer than sensations and pMB, whereas for Pattern 2, MW and pMB were significantly closer than sensations and sMB. For Pattern 3, sensations were significantly closer to this pattern than MW and pMB, and sMB was closer than pMB.

### 1.3. Self-caught MB is linked with lower physiological arousal compared to MW

To investigate physiological arousal across mental states, we analyzed the magnitude of the GS (Fig 4A). We found that sMB presented a significantly higher GS magnitude than MW episodes (MW-sMB: Δ = -0.001, 95% CI [-0.002, -0.0001], z = -3.17, *p* = .008, Fig 4B), a marker of reduced vigilance (Liu et al., 2017; Wong et al., 2013). Thus, our findings suggest that sMB reports occur during periods of lower physiological arousal relative to MW. No other significant differences were observed between the remaining mental states (*p* > .05). These results remained stable when adding the task length for each subject as a covariate (S8).

**Fig 4.**
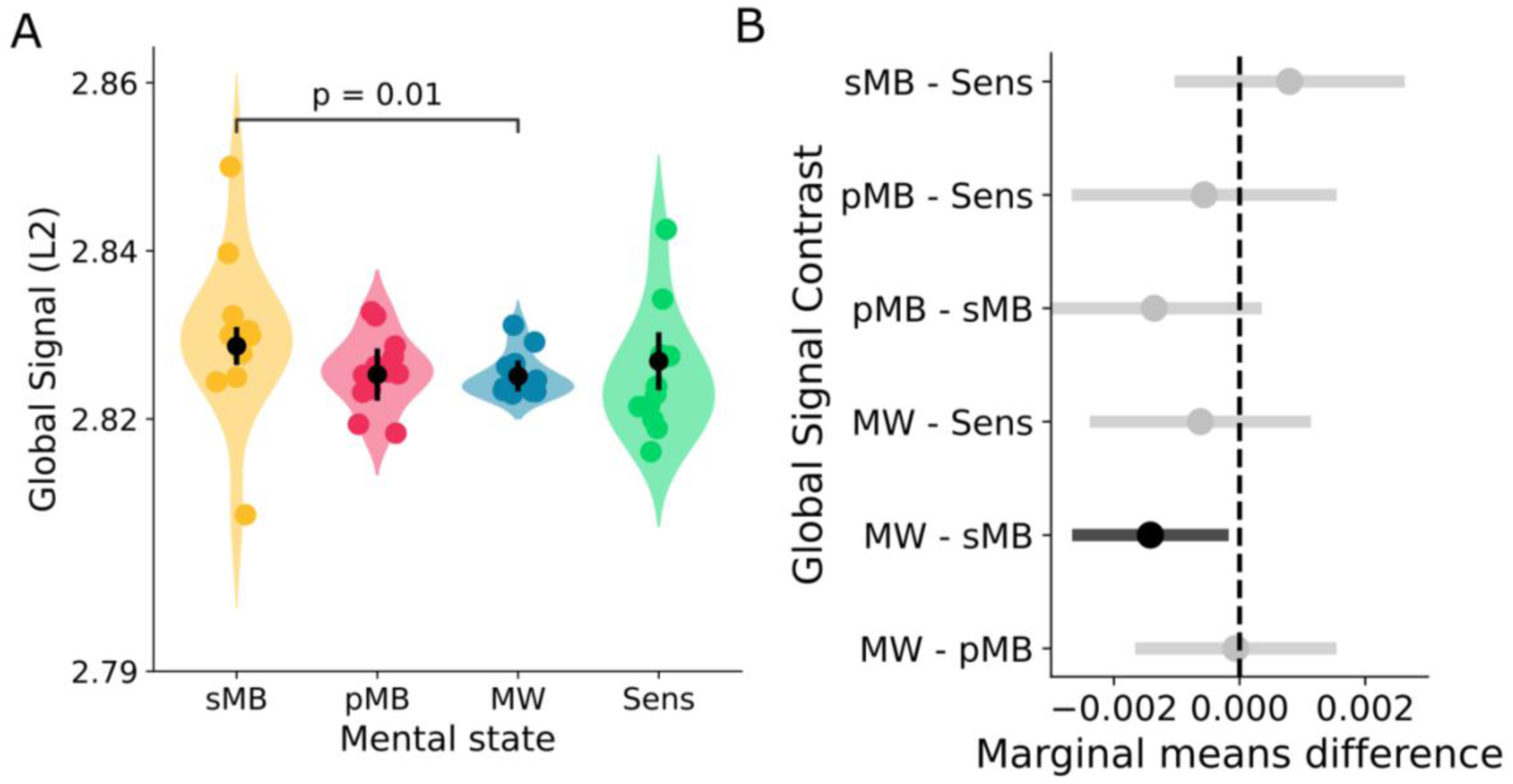
Self-caught MB reports showed higher fMRI global signal magnitude compared to MW, pointing to the potential implication of low cortical arousal when self-assessing the presence of content-less experience. A) Boxplot comparing global signal magnitude around mental states reports. Each dot represents an average of one subject’s reports. B) Statistics from marginal means contrast between the global signal characterizing different mental states reports (adjusted for multiple comparison via Tukey method) indicates that sMB has a significantly higher fMRI global signal magnitude compared to MW (Significant contrasts are black-colored). MW = mind wandering, pMB = probe-caught mind blanking, sMB = self-caught mind blanking, Sens = sensations.

### 1.4. Self-caught MB recruits greater sensorimotor/cingulate activation, and temporal, frontal and occipital deactivation compared to probe-caught MB

We compared the neural signature underlying to the two types of MB reports considering all the subjects that reported both types of MB (*n*=13). The full table of FDR-corrected voxels is available in supplementary (S9), and the unthresholded z statistical map can be explored via NeuroVault (https://neurovault.org/images/1040003/). Here, we report FDR-surviving voxels aggregating into clusters ≥ 216 mm³. This threshold approximates the size of one resolution element (also known as “resel”, see Worsley et al., 1996), calculated as the cube of the applied smoothing kernel’s FWHM (6 mm).

For positive activations, sMB showed greater recruitment, relative to pMB, in the anterior cingulate cortex (ACC), postcentral gyrus, and inferior temporal gyrus. Deactivations were observed across occipital, temporal, and frontal pole regions (Fig 5). The unthresholded z-map uploaded to NeuroVault was decoded using Neurosynth (Yarkoni et al., 2011), a meta-analytic term-based decoding tool that identifies terms whose associated activation maps are most spatially correlated with the input map. The terms most strongly associated with the sMB > pMB contrast map were predominantly related to supplementary motor and sensorimotor areas.

**Fig 5.**
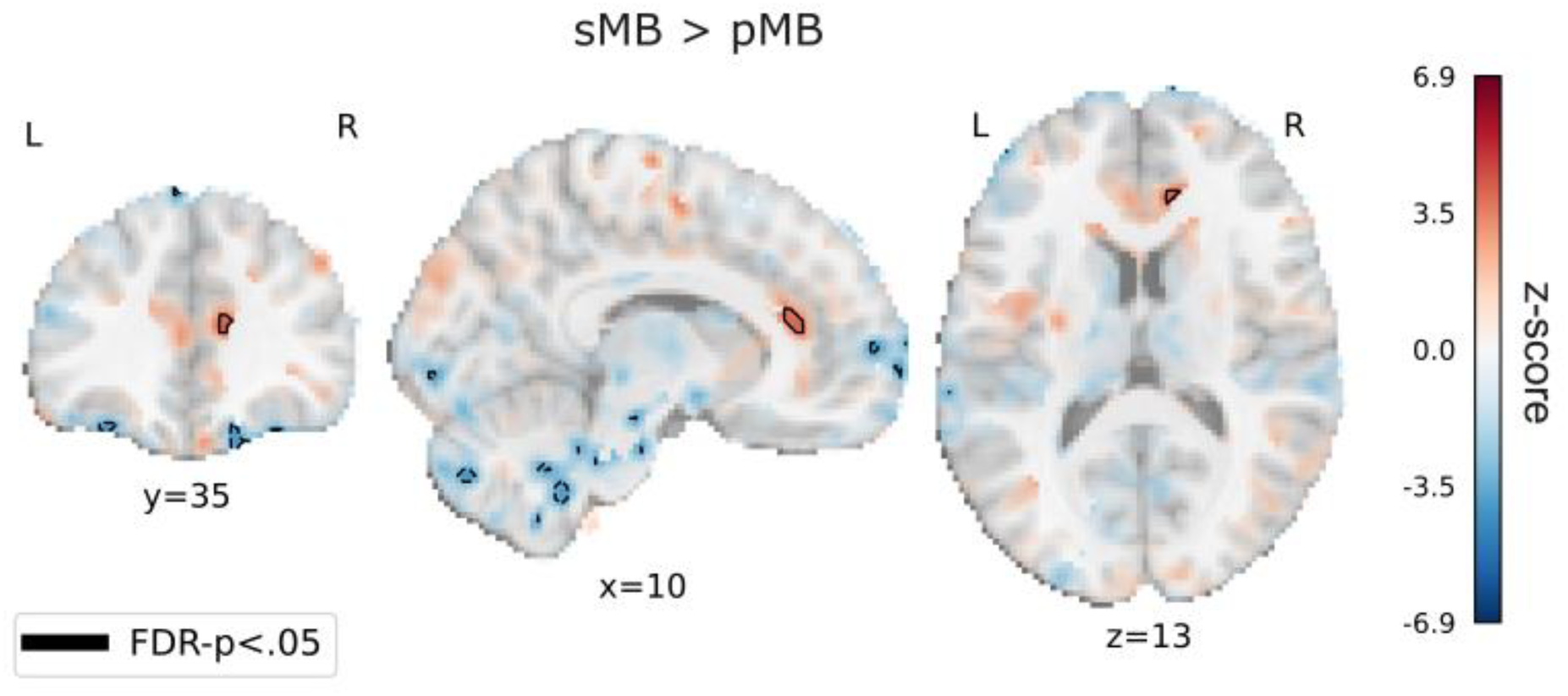
Self-caught MB recruits activations of anterior cingulate and (pre)motor regions, and greater deactivations of occipital, temporal and frontal areas. Voxel-wise fixed-effects contrast sMB>pMB shown at un uncorrected threshold pf *p*<.001. Red voxels indicate grater activations, and blue voxels indicate greater deactivations for sMB relate to pMB. Black contours outline clusters surviving FDR correction p<.05.

## 1. Discussion

We here compared two self-report methodologies for capturing MB events during typical waking time, the widely-used probe-caught method (Boulakis, Simos, et al., 2025; Koroma et al., 2023; Munoz-Musat et al., 2025; Robison et al., 2019; Unsworth et al., 2021; Van Calster et al., 2017), and the self-caught method (Ward & Wegner, 2013). Our results indicate that both methods successfully capture MB episodes, which share similar behavioral properties and are characterized by hyperconnectivity across the brain. Where these states diverged was in BOLD activation, with sMB recruiting greater ACC and (pre)motor region activation, and occipital, frontal and temporal deactivations, compared to pMB.

A primary contribution of this work is the exploration of sMB, demonstrating that sMB shares convergent neurobiological correlates with pMB. Unlike previous paradigms that required participants to intentionally “empty their minds” (Kawagoe et al., 2019), the current approach allows participants to report if they notice their mind went blank spontaneously. The significant correlation between sMB and pMB frequencies across individuals suggests high cross-method consistency, indicating that both detection methods track individual differences in the propensity to experience MB within the experimental environment. Furthermore, the uniform temporal distribution of reports in both tasks aligns with previous evidence suggesting that MB occurrence is not time-dependent, but a default mental state taking place in the stream of ongoing thinking (Mortaheb et al., 2022). While Ward and Wegner (2013) also proposed that overall MB reports appeared steadily over time, they observed a diverging trend over time where pMB reports increased while sMB decreased. It should be noted, though, that those findings were observed during a task condition (reading) and over a longer duration (20 minutes), which could have led to a lower capacity to monitor one’s stream of thought due to fatigue. In the present analysis, both tasks took place at a shorter resting-state context (8 minutes), which could have permitted the preservation of sufficient vigilance so that both detection methods led to capture the MB phenomenon stably across time.

Additionally, we show that with the probe-caught task, MB was reported in approximately 10% of probes, which is consistent with previous literature at large (for a meta-analysis on MB rates see (Boulakis et al., 2026)). However, given that the frequency of these reports is probe-dependent, episodes occurring between probes are inherently missed. While self-caught reports were more frequent in absolute count, the two methods differed in sampling rate and task duration, making a direct numerical comparison difficult: self-caught reports were collected continuously over acquisition time, whereas probe-caught reports were constrained to the probes. Nevertheless, this descriptive pattern aligns with Ward and Wegner (2013), who similarly observed that sMB events consistently outnumbered pMB episodes during a reading task. Our results point in a similar direction and extend this trend to a resting-state condition, notwithstanding the task instruction to self-detect and report these blank moments.

We consider that the higher absolute frequency of sMB relative to pMB suggests a critical methodological trade-off. While the self-caught method provides a continuous real-time reporting that can capture events that would otherwise be missed between intermittent probes, the explicit requirement for metacognitive assessment of sMB may create a dual bias. On the one side, it might redirect the participant’s focus to internal monitoring, thereby increasing false positive detection of MB (i.e., over-sensitivity). On the other side, it could as well paradoxically underestimate true frequency, because the continuous cognitive load of self-monitoring may actively prevent the mind from entering a blank state. Moreover, as self-caught methodologies by definition only capture mental states with meta-awareness (Chu et al., 2023), they necessarily exclude instances without meta-awareness. A possible alternative for future research to resolve this reporting paradox is to move towards report-independent decoding techniques. For instance, by using the neural signatures (i.e., hyperconnectivity, decrease in GS) as features in a decoder scheme, it may be possible to validate the occurrence of MB episodes independent of the participant’s active report in a more representative way. Initial efforts to decode MB from brain markers (Munoz-Musat et al., 2025), as well as from body-brain signals (Boulakis, Simos, et al., 2025), have already been proved successful. Indeed, these protocols involved training classification algorithms that relied on MB reports as the ground truth. While Boulakis et al. (2025) applied these models to predict other labeled MB reports, Munoz-Musat et al. (2025) took an initial step towards extrapolation by applying their trained classifier to unprobed, unlabeled data blocks during a cognitive task. By crossing these unlabeled predictions against behavioral performance (e.g., increased miss rates and prolonged reaction times), they provided an indirect behavioral validation of the decoded MB episodes. Ultimately, efforts to develop classification schemes free from reports could provide the ideal middle alternative: they do not confound MB with the detection itself (like self-caught methods), but also allow for a continuous tracking without being limited to discrete sampling windows (like probe-caught methods).

We also identified that both MB capturing methodologies converged on their connectome signature, associated with the hyperconnected brain state (Pattern 4), which lacks the variability in connectivity linked to typical content-full cognition (Deco et al., 2017). It is important to mention that this hyperconnected brain pattern has also been observed in states of reduced consciousness, such as during sleep and under anesthesia (Aedo-Jury et al., 2020), which reinforces the hypothesis that MB is characterized by a physiological and neural profile of modified arousal. This point is here supported by the significantly higher GS magnitude during sMB compared to content-full states, such as MW. Given that high GS amplitude is a robust marker of low cortical arousal (Liu et al., 2017; Wong et al., 2013), these findings add to previous findings that MB’s brain correlates resemble those of sleep-like activity, as previously supported by the detection of local slow wave activity during MB episodes (Andrillon et al., 2021; Munoz-Musat et al., 2025). Interestingly, the association with low arousal markers appeared more robust in the self-caught reports. This pattern may point toward two distinct mechanisms. On the one hand, it may stem from the nature of the task: as participants continuously monitor their cognition, they may more readily identify a MB episode whenever they perceive a relevant decrease in vigilance. On the other hand, the task itself may actively induce this state: the sustained cognitive effort of continuous monitoring could produce mental fatigue, hence leading to a reduction in overall arousal levels and thereby leading to reporting MB episodes. It is important to note, though, that sleep and MB are not the same state. Phenomenologically, individuals distinguish the feeling of an empty mind from the sensation of drowsiness, and reporting frequencies of MB and sleep do not correlate (Boulakis, Simos, et al., 2025). As a limitation, in our protocol, while participants were explicitly instructed to report MB, they were not offered the choice of “Sleep”, which could have facilitated a clearer interpretation of the underlying mechanism of the two MB capturing methods.

What we found, though, was that pMB reports did not significantly differ in their GS magnitude levels compared to other mental states and were also associated with a FC pattern characterized by anticorrelations between the DMN and sensory networks (Pattern 2), for which MW was also associated with. These finding may arise from the fact that probe-caught tasks may capture a boarder, more heterogeneous range of blanks (Chu et al., 2023). For instance, pMB often aggregates both the introspective experience of an empty mind and the retrospective inability to retrieve previous thoughts (Boulakis et al., 2026). This aligns with our recent study in the context of an attention task, where pMB was associated not only with the hyperconnected brain FC pattern, but also with an anticorrelation FC pattern, and this effect was mediated by perceived levels of vigilance: when people reported being alert, the connectomes around MB reports resembled the hyperconnected pattern (Boulakis, Kusztor, et al., 2025). Overall, the broad associations between pMB with distinct FC configurations may reflect the different kinds of experience and psychological mechanisms that are grouped under the umbrella term MB (Boulakis et al., 2026; Boulakis & Demertzi, 2025).

Regarding the activation analysis, we consider that the finding that sMB recruits the ACC compared to pMB likely reflects aspects of meta-awareness, meta-cognitive evaluation, and monitoring processing required for self-detection (Mazor & Fleming, 2020). Indeed, sMB requires active, continuous meta-awareness to detect a MB event where one must not only be in a MB state, but also become aware of being in that state, while pMB relies on a passive, reactive appraisal prompted by the external probe (Chu et al., 2023). As the ACC is tightly involved in metacognition (Vaccaro & Fleming, 2018) and self-referential processing (D’Argembeau et al., 2007), we propose this activation reflects the internal evaluation of their ongoing cognition. Activity on the ACC has only been observed during MB when participants are asked to empty their minds intentionally (Kawagoe et al., 2019), but no such activity has been observed during spontaneous pMB (Boulakis et al., 2023). This is also reflected in the time-varying functional connectivity pattern, as sMB was associated with a pattern of high within-somatosensory network connectivity, which was not the case for pMB. This could be tapping into distinct nuances of MB: with and without meta-awareness, as similarly observed for MW (Christoff et al., 2009; Chu et al., 2023; Seli et al., 2017).

However, this ACC and sensorimotor recruitment cannot be fully separated from a motor activity. Because MB events were defined relative to the self-initiated button press marking detection, the sMB events inherently include preparation for a self-generated motor response, even if the button press event was regressed out. This is not the case for pMB, where the mental state and the evaluation are temporally separated: the pMB event is defined prior to probe onset, while the passive evaluative process begins after probe onset until button press. Given the large recruitment of sensorimotor areas alongside the cingulate cortex in sMB, these activations likely reflect a combination both of metacognitive and motor initiation processes. On the other side, the widespread deactivations at occipital, temporal and frontal areas could be related to reduced attention to external visual or auditory stimuli, leading to the report of a MB. However, it is worth noting that this analysis was based on *n*=13 subjects and used a fixed effects approach and therefore cannot be extrapolated outside this sample.

This study presents some limitations. A primary drawback is our relatively small sample (*n*=22), although this is comparable to other MB studies (Adam et al., 2020; Torres et al., 2024). Notably, the activation analysis could only be performed on a subset of participants who reported MB episodes in both tasks, effectively reducing the sample size to *n*=13, a challenge inherent to MB low report frequency. To circumvent this limitation, we performed a fixed-effects analysis instead of mixed-effects. While this approach increases statistical power, it also prevents us from generalizing the results to the population. Furthermore, and as mentioned above, an artifact of the different methodologies is that the MB window of analysis differed slightly between tasks. For the self-caught task, MB was defined immediately preceding the button press, whereas for the probe-caught task, the mental state was defined prior to the probe appearance. This inevitably places sMB reports in closer temporal proximity to the motor response and its associated movement preparation. Even though the motor response was regressed out from the BOLD time-series, this temporal proximity could potentially contribute to the observed activity in the motor related areas. Lastly, we did not include “Sleep” among the probe-caught choices, an addition that could have aided the interpretation of our findings.

To conclude, our findings suggest that sMB resembles pMB in their behavioral properties and brain correlates of hyperconnectivity, although they diverged in the recruitment of the ACC and (pre)motor areas, a difference we attribute to the meta-cognitive process and preparation for movement initiation involved inherent to the self-caught task, alongside deactivation in sensory areas. This positions the self-caught methodology as an adequate tool to detect MB episodes. This work provides a new framework for studying MB, enabling new experimental protocols, where probe-caught approaches might be disrupting the stream of thought.

## Acknowledgements

The experimental work was conducted at the GIGA-In Vivo Imaging/Cyclotron platform of ULiège, Belgium. The article was supported by the Belgian Fund for Scientific Research (FRS-FNRS), the European Union’s Horizon 2020 Research and Innovation Marie Skłodowska-Curie RISE programme *NeuronsXnets* (grant agreement 101007926), the European Cooperation in Science and Technology COST Action “NeuralArchCon” (CA18106), the Léon Fredericq Foundation, and the University and the University Hospital of Liège.

## Data and Code Availability

Dataset is available at Dataverse https://doi.org/10.58119/ULG/EN7QJX [under review for publication] and https://gitlab.uliege.be/poc/public_datasets/self-caught-mb-fmriprep. All codes are available at https://gitlab.uliege.be/poc/self-caught-mb.

## Authors Contributions

AD, SM, and PAB conceived and designed the study. SM performed the fMRI data acquisition. AAD conducted the behavioral and neuroimaging analyses, created the figures, and wrote the original manuscript draft. Supervision was provided by AD, FR, KP, PAB, GV, and FC. Infrastructure, resources, and project administration were provided by GV and FC. All authors reviewed, edited, and approved the final version of the manuscript.

## Acknowledgements

The experimental work was conducted at the GIGA-In Vivo Imaging/Cyclotron platform of ULiège, Belgium.

