## Supplementary Material for "How to catch a blank mind? Brain similarities and differences between self- and probe-caught mind blanking"

For

### Supplementary 1: Frequency of other mental states during the probe-caught task.

The Table 1 summarizes the mental state frequency (over the total N probes) during the probe-caught task.

***Table 1.*** *Frequency of mental states.*

| Comparison | Frequency |
|---|---|
| Mind Wandering | 73.65% |
| Sensation | 11.52% |
| Mind Blanking | 10.8% |

### Supplementary 2: Baseline comparisons between resting connectivity profiles in the probe-caught and self-caught task.

This analysis checks for baseline differences in pre-defined rest functional connectivity (FC, baseline periods without mental state reports) between the probe-caught and self-caught tasks, which could bias k-means clustering or distance metrics in the main analysis.

First, we expected that rest FC matrices from both tasks would form similar clusters if no systematic differences existed. By applying the same k=4 clustering separately to rest FC matrices from each task, we show no differences in connectivity patterns of the rest periods of either method. Figure 1 displays the resulting patterns, with the third row showing residuals (i.e., difference) between corresponding self-caught and probe-caught clusters.

To quantitatively assess that the centroids do not differ, we also computed cosine distances for each rest FC matrix to the four centroids from its own task and the opposite task (Figure 2). We then fitted Bayesian linear mixed-effects models (via brms) to test the fixed effect of task (response: self-caught vs. probe-caught) on scaled distances to each cluster centroid, including random intercepts and slopes by subject. Equivalence was assessed using Bayesian equivalence testing with a ROPE of ±0.455 (medium-to-large standardized mean difference threshold, given the large sample of FC matrices). Most clusters showed equivalence (ROPE % = 1, HDI within ±0.455), supporting similar rest baselines across tasks, with two undecided cases (Table 2).

***Figure 1***. *Connectivity clusters calculated on the rest periods for each task separately indicated no differences, thus ensuring similar baseline states.*

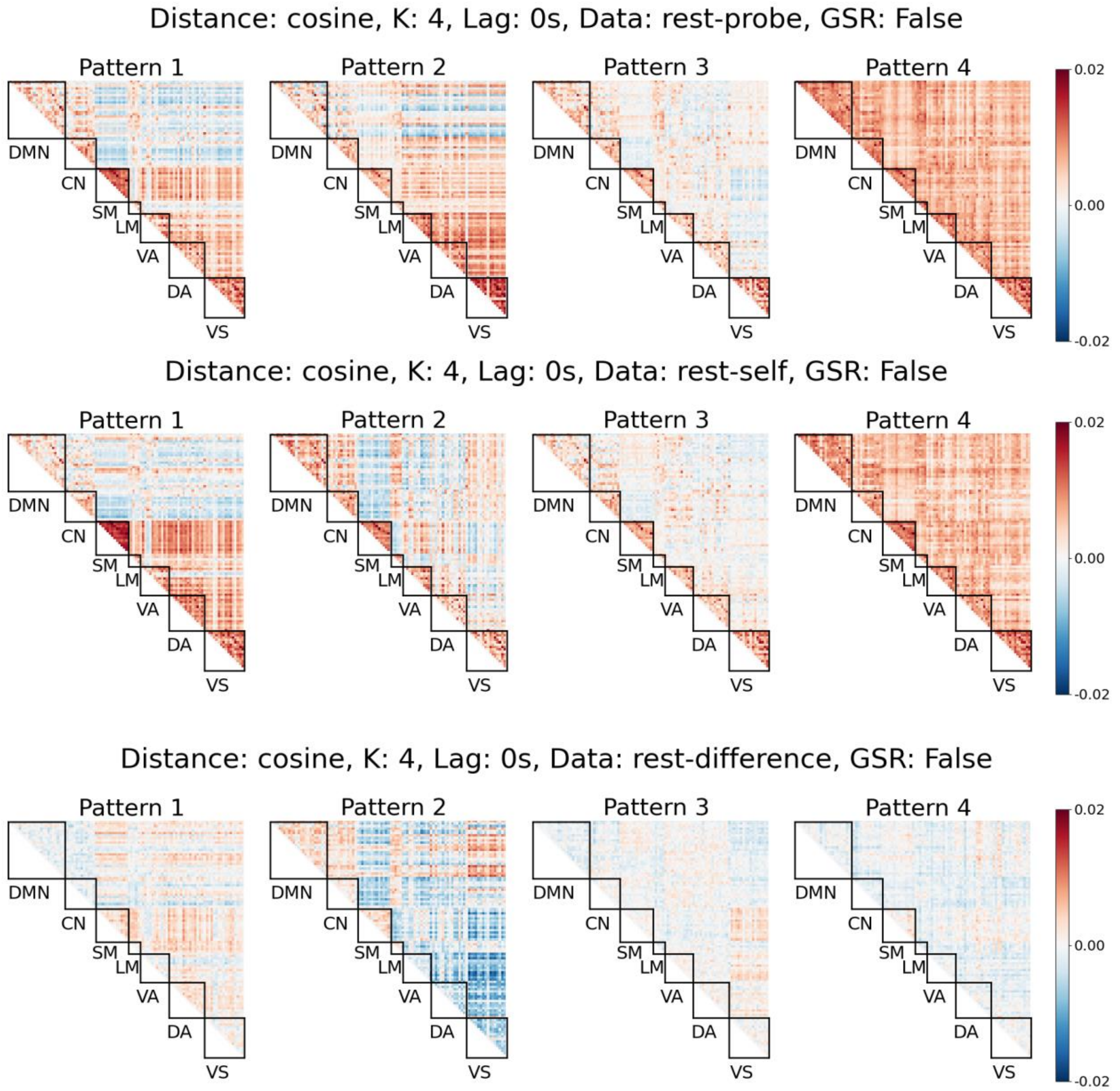


Resulting clusters from probe-caught resting data (upper) and self-caught resting data (middle) and their difference (lower). Patterns of probe-caught are organized by metastability. As self-caught patterns organized by metastability did not match the probe-caught, they were visually aligned to calculate the differences.

***Figure 2.*** *Distance of each functional connectivity matrix to the centroids of its own task and the opposite task show equivalence (Bayesian Test)*

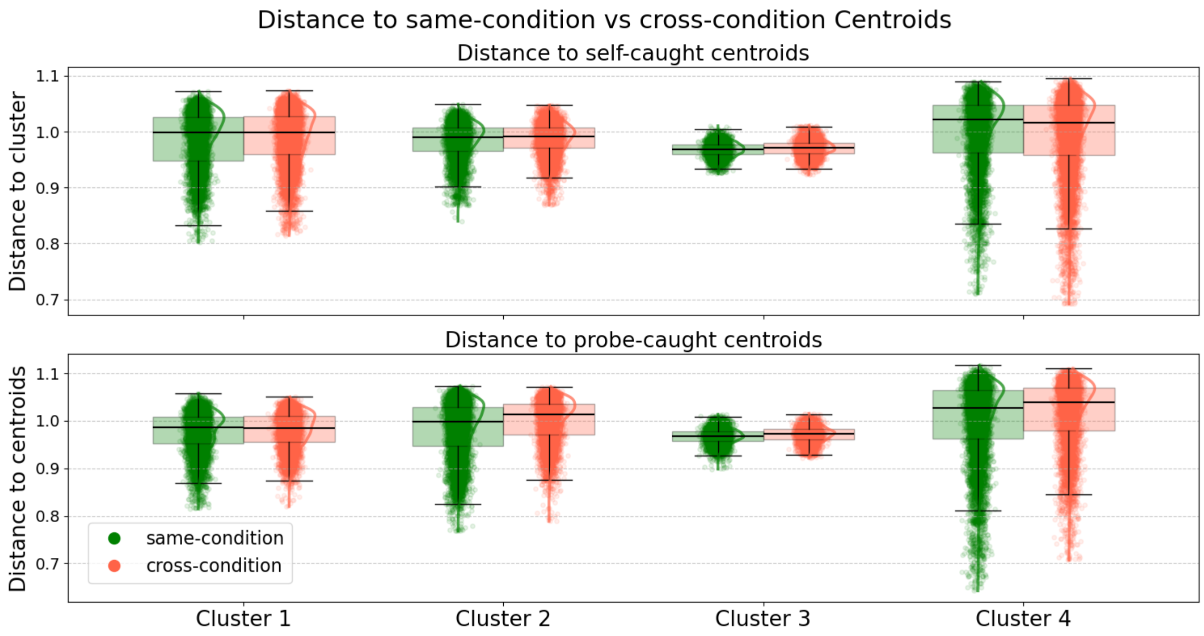

Distances of each FC matrices to its own task centroids (i.e, upper plot shows self-caught FC distances to self-caught centroids in green and probe-caught FC distances to self-caught centroids in orange; otherwise for the lower plot showing probe-caught data).

***Table 2.*** *Comparison of functional connectivity matrix to the same- and cross-task rest centroids*

| Comparison | HDI low | HDI high | ROPE % | Equivalence Decision |
|---|---|---|---|---|
| Self-Caught Cluster 1 | -.41 | .12 | 1 | Accepted |
| Self-Caught Cluster 2 | -.36 | .16 | 1 | Accepted |
| Self-Caught Cluster 3 | -.51 | -.05 | 95 | Undecided |
| Self-Caught Cluster 4 | -.18 | .15 | 1 | Accepted |
| Probe-Caught Cluster 1 | -.22 | .26 | 1 | Accepted |
| Probe-Caught Cluster 2 | .09 | .47 | 99 | Undecided |
| Probe-Caught Cluster 3 | .06 | .45 | 1 | Accepted |
| Probe-Caught Cluster 4 | -.05 | .24 | 1 | Accepted |

HDI = Highest Density Interval; ROPE = Region of Practical Equivalence.

### Supplementary 3: Optimal k parameter for clustering

To determine the optimal number of clusters (k), we evaluated several internal validation metrics, as different indices prioritize different structural properties of the data. First, we calculated heuristic and geometrics measures such as Inertia (sum of squared distances to the nearest cluster center; also known as Elbow Method) and the Silhouette Coefficient, which assesses cluster separation and cohesion. Second, we employed variance-ration indices such as the Calinski-Harabasz Index and the Davies-Bouldin Index, which evaluate the ratio of within-cluster dispersion to between-cluster separation. Lastly, we assessed the inter-pattern similarity by correlating the FC cluster centroids with one another. For each value of k, we computed the pairwise Pearson correlation between centroids and calculated the mean of mean correlation values (lower value reflects less correlation across centroids).

As illustrated in Figure 3, the results did not converge on a single optimal k.

While the metric silhouette score suggested k=6, the Calinski-Harabasz index pointed toward k=3, and the Davies-Bouldin index suggested k=3 and k=4. The Inertia method did not point towards a clear optimal k, and the mean Inter-pattern correlation pointed towards k=6. In the absence of a clear convergence across indices, we adopted a literature-driven approach. We proceeded with k=4, as four stable resting-state patterns have been consistently identified in previous functional imaging studies (Demertzi et al., 2019; Mortaheb et al., 2022). This choice ensures both biological plausibility and direct comparability with existing research.

***Figure 3***. *Internal validation metrics do not converge on a single optimal k*

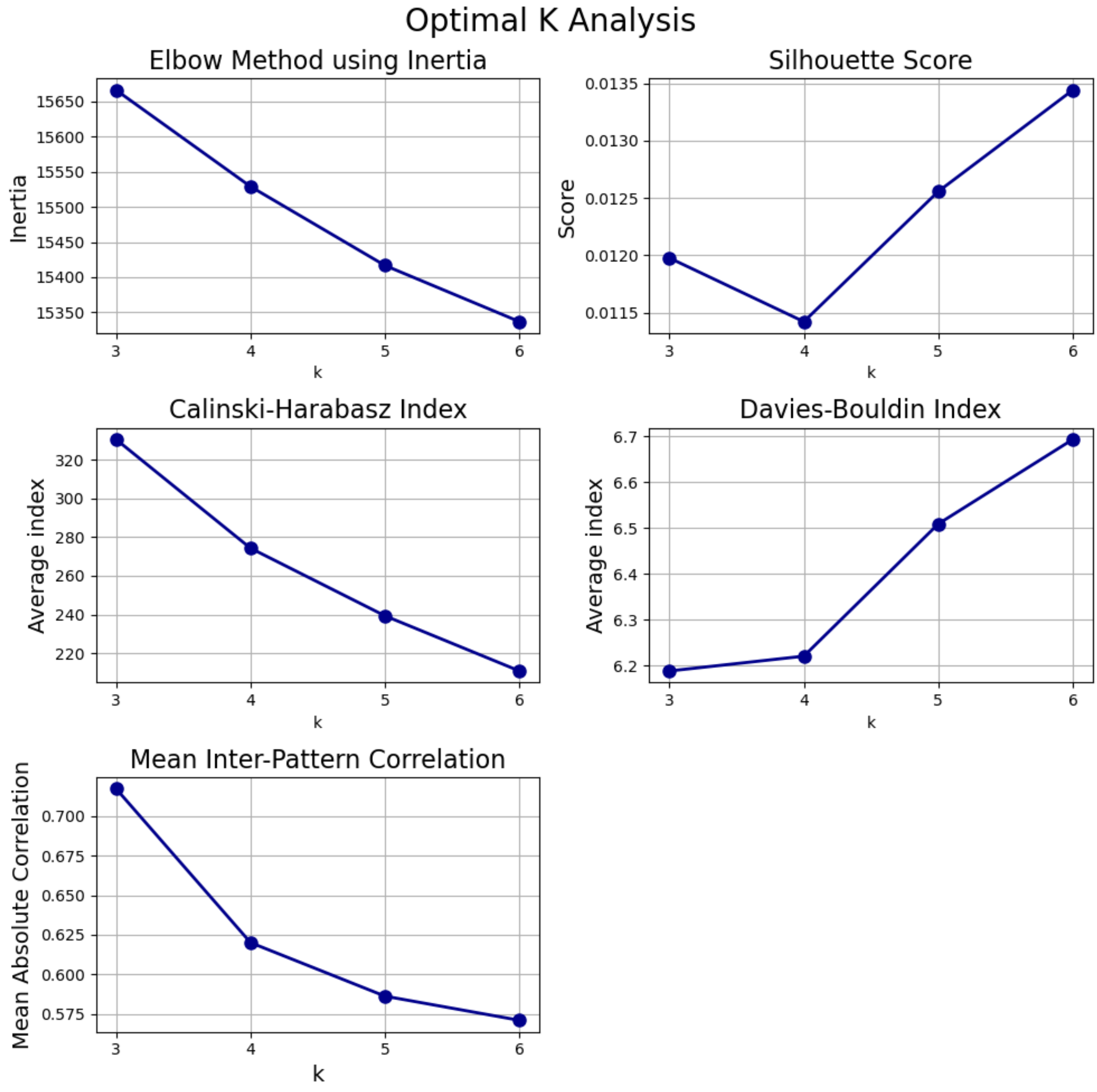


Each panel shows a different internal validation metric used to assess the optimal number of clusters (k) for k-mean clustering across k=3 to k=6.

### Supplementary 4: Consistency of patterns and distance analysis with varying parameters

To assess the stability of the identified patterns across different clustering solutions with varying *k*, we performed a cross-k centroid stability analysis. Given that centroids were ordered by metastability, this analysis was used to determine whether the ordered centroids remained consistent when the number of clusters (k) varied. Using the k=4 as a reference, we correlated its first and last centroids with the centroids obtained at other values of k. This approach showed that the last centroid was highly stable across all tested values of k, whereas the first centroid remained stable at lower k, but shifted in ordered position starting at k=5.

The change in centroid position starting at k=5 suggests that one of the patterns may be subdivided into more than one related cluster as the solution becomes finer. This supports the selection of k=4 as a parsimonious solution, because it captures the main pattern structure while avoiding unnecessary fragmentation of similar states. Overall, these results support the reproducibility of the main patterns across cluster solutions. Actual centroids for varying k solutions can be observed in Figure 5, including a summary visualization with PCA (n components =2).

***Figure 4.*** *Cross-k centroid Matching with anchor in k=4*

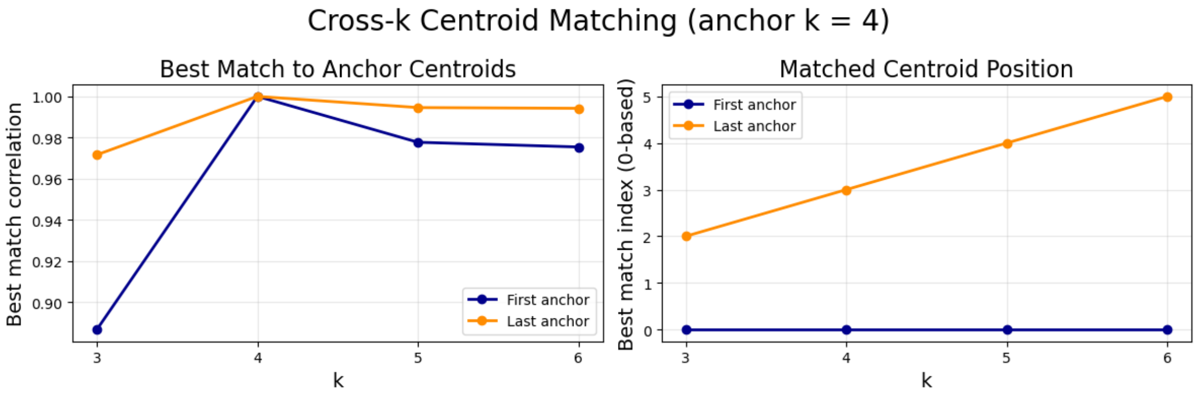


The left panel shows the correlation between the first centroid (blue) and last centroid (orange) across different values of k, using the k=4 as the reference anchor. The right panel shows which cluster had the highest correlation with the first centroid of the k=4 clustering (blue) and last centroid of the k=4 anchor (orange). The expected result is that the first anchor centroid matches the first ordered centroid in the other k-means clustering, and the last anchor centroid matches the last ordered centroid.

***Figure 5.*** *K-means clustering with varying k*

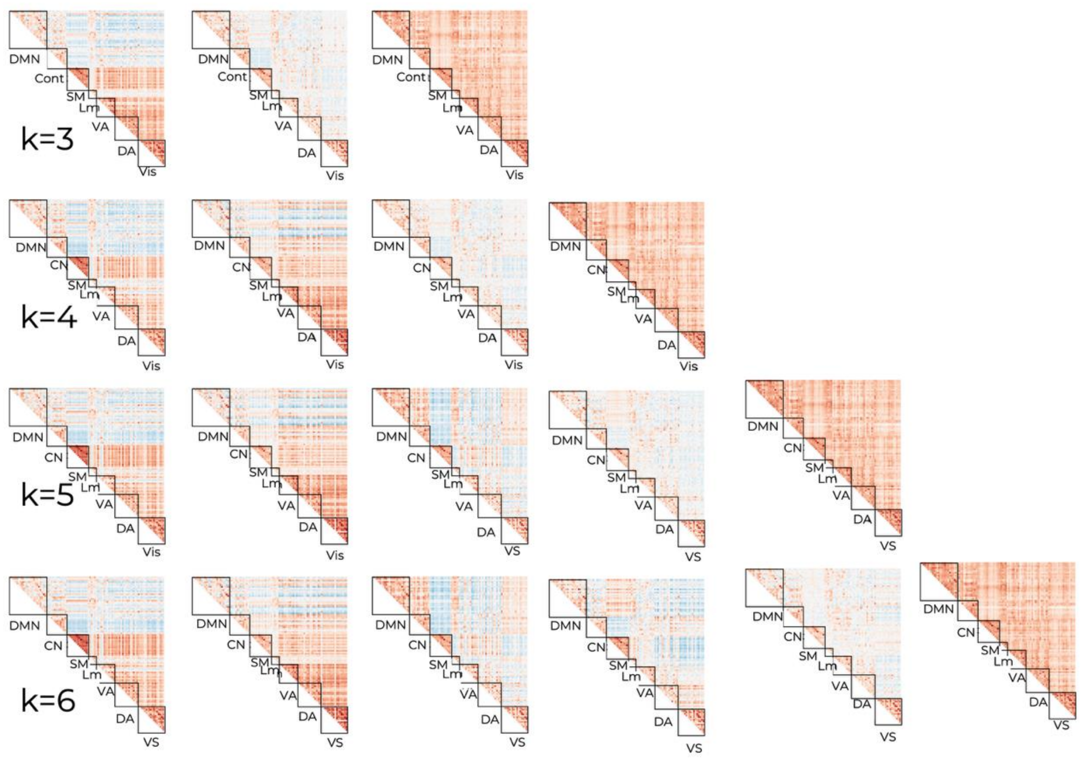


Centroids for k-means clustering varying from k=3 to k=6, ordered by metastability.

Mental states FC matrices distance to centroids remained consistent from k=3 to k=5. While some neuro-coupling results vary, the relation between MB and the hyperconnected pattern (our primary focus) stays stable across solutions (Figure 6).

***Figure 6.*** *Distances of each mental state FC to the centroids of k=3 and k=5 replicate results of k=4*

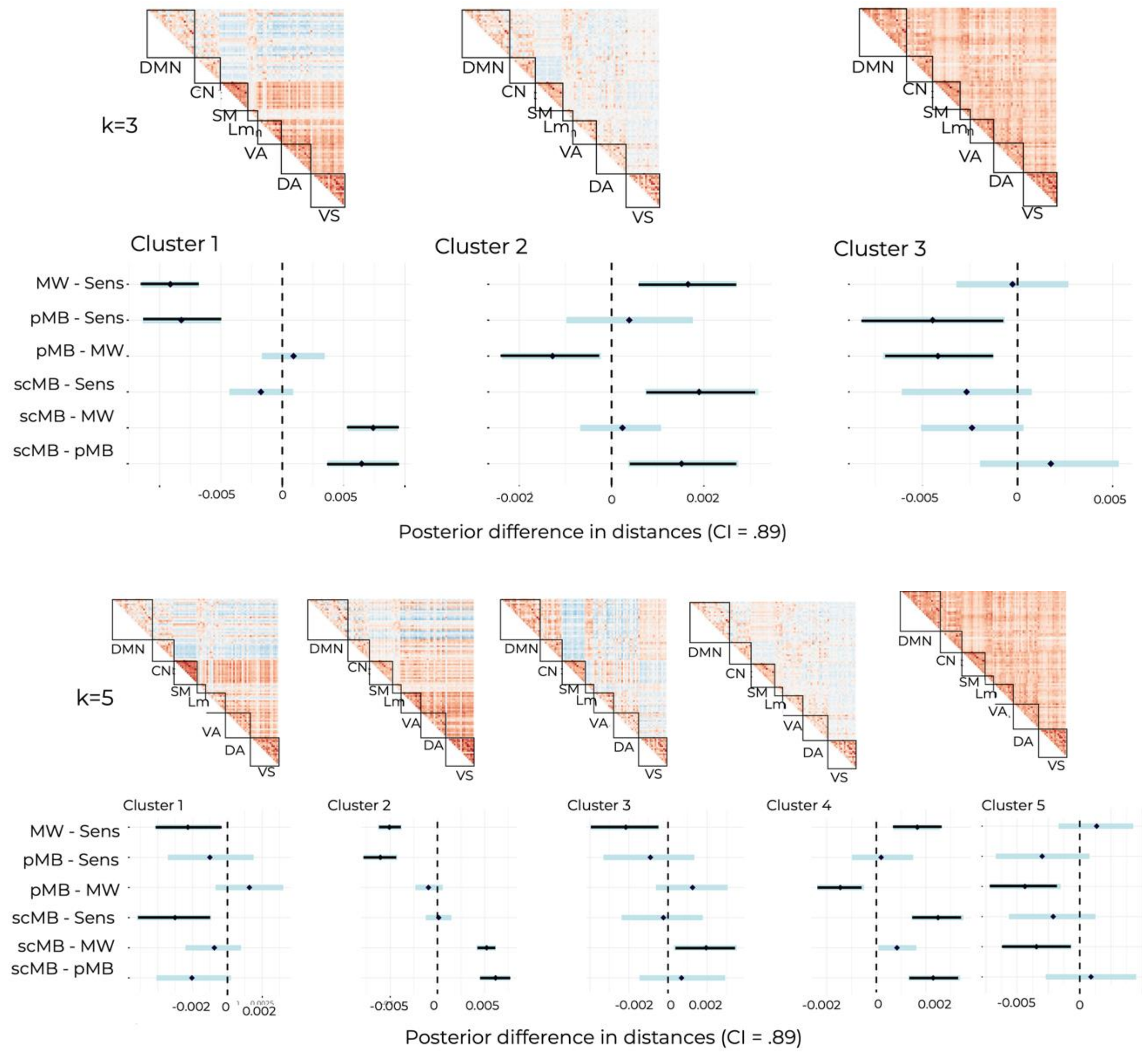


K-means centroids of time-varying functional connectivity and connectome. Scale shows Fisher Z and L2 normalized FC. Estimated Marginal Means contrast between distances of the mental state functional connectivity matrices to each centroid. Significant contrasts are red-colored.

### Supplementary 5: Occurrence rate of functional connectivity patterns

The occurrence rate of each pattern was estimated counting the number of matrices assigned to each centroid for each participant. We compared occurrence rates with a paired t-test with $p$-FDR < 0.05 corrected for multiple comparisons. Figure 7 shows the occurrence rate of the functional connectivity centroids. Pattern 3 presented a significant higher occurrence than all other patterns).

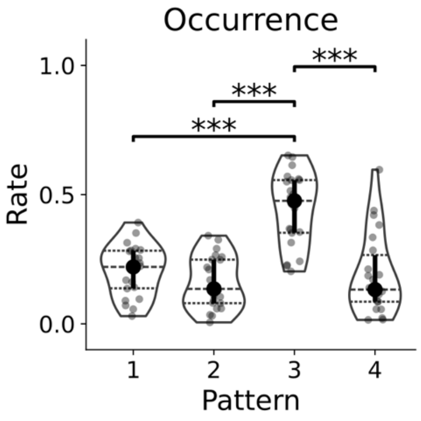

***p<.001.

### Supplementary 6: Comparison Null vs Response Model

Table 3 shows the Bayes Factor, Akaike Information Criterion (AIC) and Bayesian Information Criterion (BIC) for the comparison between the null model and response model. Null model explains distances to that cluster just with the intercept. Response model included mental state reports as fixed effects and subject-specific random intercepts.

Results were mixed across measures and models. Bayes factors favored the null model in Clusters 1, 3, and 4, but favored the response model in Cluster 2. BIC favored the response model for Clusters 3 and 4; and the AIC for Cluster 3.

***Table 3.*** *Comparison null and response model for each cluster.*

| Model | BF | AIC | BIC |
|---|---|---|---|
| **Response Cluster 1** | -14.41 | 1.036 | -1.798 |
| **Null Cluster 1** | | 0.460 | -2.308 |
| **Response Cluster 2** | 20.27 | -0.011 | -2.683 |
| **Null Cluster 2** | | -0.016 | -2.695 |
| **Response Cluster 3** | -16.082 | 0.060 | -2.622 |
| **Null Cluster 3** | | 0.273 | -2.492 |
| **Response Cluster 4** | -14.175 | 0.009 | -2.553 |
| **Null Cluster 4** | | 0.005 | -2.531 |

BF = bayes factor, AIC = Akaike Information Criterion, BIC = Bayesian Information Criterion.

### Supplementary 7: Proximity of Mental States to Functional Connectivity Patterns

Table 5 shows post-hoc comparisons of distances between mental states functional connectivity and functional connectivity centroids. Contrasts with 89% credible intervals excluding zero are considered supported (Bonferroni-corrected threshold for six pairwise comparisons).

***Table 5.*** *Estimated marginal means between mental states distances to k-means clusters*

| | Contrast | Δ | 98% CI | |
|---|---|---|---|---|
| **Pattern 1** | **scMB - pMB** | -0.003 | -0.0062 | -0.0014 |
| | **scMB - MW** | -0.0004 | -0.002 | 0.0014 |
| | **scMB - Sens** | -0.002 | -0.005 | -0.0005 |
| | **pMB - MW** | 0.0033 | 0.0011 | 0.0054 |
| | **pMB - Sens** | 0.0008 | -0.0019 | 0.0035 |
| | **MW - Sens** | -0.0024 | -0.0044 | -0.0002 |
| **Pattern 2** | **scMB - pMB** | 0.01145 | 0.0083 | 0.0146 |
| | **scMB - MW** | 0.009 | 0.0074 | 0.0114 |
| | **scMB - Sens** | 0.000 | -0.0026 | 0.0028 |
| | **pMB - MW** | -0.001 | -0.0047 | 0.0010 |
| | **pMB - Sens** | -0.011 | -0.0149 | -0.0078 |
| | **MW - Sens** | -0.009 | -0.0119 | -0.0069 |
| **Pattern 3** | **scMB - pMB** | 0.0016 | 0.0004 | 0.0028 |
| | **scMB - MW** | 0.0007 | -0.0001 | 0.0015 |
| | **scMB - Sens** | 0.0022 | 0.0010 | 0.0035 |
| | **pMB - MW** | -0.0009 | -0.0019 | 0.0001 |
| | **pMB - Sens** | 0.0006 | -0.0007 | 0.0020 |
| | **MW - Sens** | 0.0015 | 0.0005 | 0.0026 |
| **Pattern 4** | **scMB - pMB** | 0.0013 | -0.0024 | 0.0049 |
| | **scMB - MW** | -0.003 | -0.0060 | -0.0005 |
| | **scMB - Sens** | -0.0023 | -0.0058 | 0.0011 |
| | **pMB - MW** | -0.0045 | -0.007 | -0.0016 |
| | **pMB - Sens** | -0.0037 | -0.007 | 0.000 |
| | **MW - Sens** | 0.0008 | -0.002 | 0.0038 |

### Supplementary 8: Global signal analysis with task-duration as covariate

We considered the possibility that global signal magnitude might vary with the duration of the fMRI session. To address this, we repeated the analysis while including task length for each participant as a covariate. The results remained the same: self-caught mind blanking showed higher global signal magnitude than mind-wandering ($p$=0.008).

***Table 4.*** *Estimated marginal means between mental states fMRI global signal magnitude*

| Contrast | Estimate | SE | LCL | UCL | Z | *p* |
|---|---|---|---|---|---|---|
| **MW - pMB** | 0.0000 | 0.0005 | -0.001 | 0.0014 | -0.10 | 0.999 |
| **MW - scMB** | -0.0014 | 0.0004 | -0.002 | -0.0002 | -3.17 | 0.008 |
| **MW - Sens** | -0.0006 | 0.0006 | -0.002 | 0.0010 | -0.96 | 0.7699 |
| **pMB - scMB** | -0.0013 | 0.0006 | -0.002 | 0.0002 | -2.16 | 0.1336 |
| **pMB - Sens** | -0.0005 | 0.0007 | -0.002 | 0.0014 | -0.71 | 0.889 |
| **scMB - Sens** | 0.0007 | 0.0006 | -0.0009 | 0.0025 | 1.17 | 0.6403 |

SE = Standard Error; LCL = Lower Confidence Limit; UCL = Upper Confidence Limit.

### Supplementary 9: Voxel-wise activation differences between MB report types

Table 3 shows the FDR-corrected voxels from the contrast sMB > pMB. The unthresholded z statistical map can be found in via NeuroVault (https://neurovault.org/images/1040003/).

***Table 3.*** *FDR-corrected voxels from contrast sMB > pMB*

| X | Y | Z | Peak Stat | Cluster Size (mm3) |
|---|---|---|---|---|
| 1.5 | 27.5 | -30.5 | 4.8089834028284395 | 216 |
| 23.5 | -36.5 | 77.5 | 4.808744556992659 | 136 |
| 47.5 | -38.5 | -26.5 | 4.7896125282787505 | 248 |
| 23.5 | 51.5 | -4.5 | 4.4600755991818 | 16 |
| 23.5 | -60.5 | -56.5 | 4.454236778968444 | 128 |
| 33.5 | -32.5 | 39.5 | 4.441096599753524 | 496 |
| -34.5 | -32.5 | 47.5 | 4.408617736916485 | 328 |
| -38.5 | -62.5 | 9.5 | 4.285885746915026 | 24 |
| 11.5 | 35.5 | 13.5 | 4.212293884124937 | 216 |
| 23.5 | 43.5 | -10.5 | 4.027100028198911 | 8 |
| 13.5 | 39.5 | -4.5 | 3.9821013548821456 | 16 |
| 17.5 | -44.5 | 37.5 | 3.9421459330155724 | 16 |
| 41.5 | -50.5 | -52.5 | 3.9218453746847977 | 96 |
| -34.5 | -42.5 | -46.5 | 3.758766324183247 | 32 |
| 23.5 | 61.5 | -16.5 | 3.754477138233353 | 32 |
| -14.5 | -72.5 | -58.5 | 3.7473067953838792 | 16 |

| | | | | |
|---|---|---|---|---|
| 41.5 | 47.5 | -10.5 | 3.7102409905589653 | 80 |
| -22.5 | -48.5 | 75.5 | 3.615997033878408 | 64 |
| -40.5 | -62.5 | 7.5 | 3.578911182436072 | 8 |
| -26.5 | 51.5 | -6.5 | 3.576419994479628 | 8 |
| 19.5 | -4.5 | 53.5 | 3.5277390255554932 | 8 |
| 23.5 | 53.5 | -6.5 | 3.5067646651333515 | 24 |
| -14.5 | 31.5 | 21.5 | 3.498973344601813 | 8 |
| -28.5 | -20.5 | 65.5 | 3.492001845332096 | 8 |
| -24.5 | 53.5 | -6.5 | 3.4355993697681013 | 16 |
| -38.5 | -34.5 | 69.5 | 3.40176504240466 | 8 |
| 17.5 | 23.5 | -6.5 | 3.3976679674290304 | 8 |
| -40.5 | -78.5 | 7.5 | 3.392573531250091 | 8 |
| -20.5 | 51.5 | -20.5 | 3.3914722773300596 | 8 |
| 11.5 | -6.5 | 57.5 | 3.386386192391602 | 8 |
| 17.5 | -90.5 | 23.5 | 3.369363457580174 | 8 |
| -28.5 | -20.5 | 69.5 | 3.3646316609794993 | 8 |
| 15.5 | 21.5 | -8.5 | 3.357200116521742 | 8 |
| 51.5 | -32.5 | 7.5 | 3.3173292319126135 | 8 |
| 39.5 | -42.5 | 67.5 | 3.3106149644827445 | 8 |
| -16.5 | -24.5 | 65.5 | 3.3068181999138204 | 8 |
| -14.5 | -40.5 | 75.5 | 3.296842303044867 | 8 |
| 19.5 | 29.5 | 9.5 | 3.295732809774177 | 8 |
| -66.5 | 3.5 | 19.5 | 3.284856193468259 | 8 |
| -34.5 | -98.5 | -6.5 | -6.910652342816279 | 3464 |
| 31.5 | 7.5 | -32.5 | -5.95190001096838 | 3480 |
| -30.5 | 29.5 | -22.5 | -5.489978914510894 | 1072 |
| -68.5 | -28.5 | 1.5 | -5.264512586912536 | 2048 |
| -62.5 | -14.5 | -34.5 | -5.243472860146549 | 448 |
| -52.5 | 5.5 | 49.5 | -5.219011794753861 | 328 |
| -56.5 | 7.5 | -34.5 | -5.126987557656519 | 4056 |
| 15.5 | -90.5 | -2.5 | -5.079203166891137 | 4896 |
| -38.5 | -14.5 | -28.5 | -5.010741593792787 | 1032 |
| -64.5 | -0.5 | -8.5 | -4.93342593871024 | 456 |
| -26.5 | -82.5 | -52.5 | -4.922146304830839 | 208 |
| -8.5 | -32.5 | -38.5 | -4.8835477580204945 | 176 |

| | | | | |
|---|---|---|---|---|
| 15.5 | -0.5 | -10.5 | -4.877316460000329 | 360 |
| 23.5 | -62.5 | 71.5 | -4.851713506302915 | 80 |
| -32.5 | -72.5 | 59.5 | -4.823424808307182 | 264 |
| 11.5 | 71.5 | -0.5 | -4.774998851665841 | 288 |
| 51.5 | 21.5 | -22.5 | -4.64153202051552 | 88 |
| 11.5 | 61.5 | 7.5 | -4.535566408211229 | 152 |
| -18.5 | 71.5 | -8.5 | -4.526686694511551 | 648 |
| 35.5 | 49.5 | -20.5 | -4.447272067126947 | 104 |
| 67.5 | -46.5 | -18.5 | -4.4454096398908325 | 56 |
| -20.5 | -40.5 | -42.5 | -4.432607590852964 | 168 |
| 15.5 | 33.5 | -26.5 | -4.4176460299705855 | 240 |
| -32.5 | 7.5 | 55.5 | -4.390642388492001 | 48 |
| 13.5 | -76.5 | -40.5 | -4.380712666255731 | 208 |
| -6.5 | 31.5 | 63.5 | -4.328292301194708 | 112 |
| 11.5 | -46.5 | -44.5 | -4.327072545030759 | 320 |
| -42.5 | -0.5 | 63.5 | -4.305685018132758 | 200 |
| -64.5 | -60.5 | -10.5 | -4.289550437111909 | 144 |
| 71.5 | -14.5 | -14.5 | -4.286650437591126 | 80 |
| 7.5 | -20.5 | -18.5 | -4.239507898125164 | 168 |
| -48.5 | -6.5 | -36.5 | -4.2316583095608085 | 536 |
| -26.5 | 29.5 | 59.5 | -4.223449956694686 | 96 |
| -68.5 | -32.5 | -12.5 | -4.202713605526534 | 528 |
| 25.5 | -16.5 | -8.5 | -4.1528688863025645 | 200 |
| 29.5 | 33.5 | -22.5 | -4.145452565759784 | 136 |
| -52.5 | 19.5 | -20.5 | -4.130783102454317 | 248 |
| 45.5 | -2.5 | -42.5 | -4.130356294144227 | 192 |
| 61.5 | -2.5 | -32.5 | -4.073028636982513 | 56 |
| -34.5 | -62.5 | -62.5 | -4.029965660341225 | 112 |
| -38.5 | -64.5 | -50.5 | -4.005131118602024 | 72 |
| 3.5 | 19.5 | -10.5 | -4.003548446484556 | 112 |
| 7.5 | -18.5 | -28.5 | -3.957451022882541 | 40 |
| -26.5 | 7.5 | -44.5 | -3.950330224826244 | 136 |
| -16.5 | -22.5 | -6.5 | -3.9426736415896144 | 64 |
| -62.5 | -18.5 | -24.5 | -3.9294380173531027 | 128 |
| -10.5 | -82.5 | -42.5 | -3.8984084461153063 | 64 |

| | | | | |
|---|---|---|---|---|
| 1.5 | -46.5 | -68.5 | -3.8979123390364823 | 32 |
| 51.5 | -0.5 | 55.5 | -3.8768060832808438 | 32 |
| -62.5 | -48.5 | 47.5 | -3.854375969876669 | 32 |
| 5.5 | -34.5 | -32.5 | -3.8417525087594058 | 168 |
| 11.5 | -50.5 | -36.5 | -3.8406635623395764 | 72 |
| -34.5 | -76.5 | -56.5 | -3.7922681153437776 | 48 |
| -46.5 | -86.5 | -6.5 | -3.785386854353831 | 120 |
| -64.5 | -34.5 | -28.5 | -3.7783197817873697 | 8 |
| -38.5 | 1.5 | 65.5 | -3.77031035470029 | 32 |
| -52.5 | -64.5 | -26.5 | -3.7650061188166326 | 104 |
| -22.5 | -60.5 | 73.5 | -3.7436546944025486 | 16 |
| -20.5 | -6.5 | -22.5 | -3.7088875399573213 | 32 |
| 17.5 | -38.5 | -10.5 | -3.6999980334552505 | 40 |
| 5.5 | -28.5 | -32.5 | -3.6987215514467326 | 32 |
| 49.5 | -0.5 | 57.5 | -3.6929392560553898 | 16 |
| 29.5 | -50.5 | -60.5 | -3.692102116428059 | 16 |
| -46.5 | 29.5 | 27.5 | -3.6912229385150717 | 80 |
| 23.5 | -78.5 | -46.5 | -3.6890499849835927 | 232 |
| -0.5 | -60.5 | -34.5 | -3.6836659310498874 | 32 |
| 23.5 | -20.5 | -8.5 | -3.6780398570938275 | 8 |
| -10.5 | -48.5 | -34.5 | -3.6735544110786984 | 56 |
| 1.5 | -28.5 | -34.5 | -3.6517189580423164 | 16 |
| -8.5 | 25.5 | -18.5 | -3.6510558353183566 | 32 |
| -54.5 | -18.5 | -16.5 | -3.6437558613092285 | 32 |
| 5.5 | 51.5 | 49.5 | -3.6371926215675177 | 8 |
| -22.5 | -28.5 | -2.5 | -3.6363803923990363 | 96 |
| 15.5 | -20.5 | -10.5 | -3.632690757521976 | 24 |
| 45.5 | -26.5 | -24.5 | -3.6274021614134284 | 8 |
| 59.5 | -18.5 | -28.5 | -3.6241023955976694 | 88 |
| -24.5 | 27.5 | 61.5 | -3.613379371821548 | 8 |
| 7.5 | 69.5 | -18.5 | -3.606911500246904 | 16 |
| -46.5 | 19.5 | 25.5 | -3.6059532407356487 | 144 |
| -6.5 | -12.5 | 3.5 | -3.5950274077235433 | 24 |
| -60.5 | -64.5 | -14.5 | -3.5753863703931694 | 32 |
| -26.5 | -12.5 | -10.5 | -3.570825851297447 | 24 |

| | | | | |
|---|---|---|---|---|
| -66.5 | -54.5 | 19.5 | -3.556423668100673 | 40 |
| -62.5 | -36.5 | -28.5 | -3.5461504695552457 | 16 |
| -46.5 | 51.5 | 15.5 | -3.5336939535777967 | 16 |
| -54.5 | -10.5 | -4.5 | -3.5312443033247605 | 40 |
| 9.5 | -54.5 | -54.5 | -3.526212470431147 | 24 |
| -14.5 | 61.5 | -20.5 | -3.524719209681297 | 32 |
| 57.5 | 13.5 | -22.5 | -3.523723764371052 | 8 |
| -64.5 | -36.5 | -26.5 | -3.507562678235079 | 16 |
| 13.5 | 27.5 | -16.5 | -3.5005272335791346 | 8 |
| 45.5 | -8.5 | -2.5 | -3.471343671050898 | 32 |
| -44.5 | -8.5 | -34.5 | -3.4682901609765606 | 8 |
| 37.5 | -2.5 | -42.5 | -3.454420953083738 | 40 |
| 25.5 | 33.5 | -24.5 | -3.450038751160353 | 8 |
| -10.5 | -58.5 | -40.5 | -3.444970133110663 | 8 |
| 15.5 | -70.5 | -28.5 | -3.4233161684795985 | 16 |
| -10.5 | -54.5 | -44.5 | -3.4177874498971956 | 16 |
| -10.5 | -54.5 | 77.5 | -3.402157430603513 | 8 |
| -34.5 | -74.5 | -58.5 | -3.3996159405865827 | 8 |
| 9.5 | -40.5 | -28.5 | -3.392997763770712 | 16 |
| 67.5 | -38.5 | -8.5 | -3.3923206722496815 | 16 |
| 15.5 | 63.5 | 5.5 | -3.3899481191032828 | 8 |
| 41.5 | -72.5 | -50.5 | -3.384288456320414 | 8 |
| 1.5 | -32.5 | -34.5 | -3.381714102769389 | 8 |
| 11.5 | 73.5 | 13.5 | -3.378742010682026 | 8 |
| -34.5 | 5.5 | 55.5 | -3.376151604648268 | 8 |
| -10.5 | -56.5 | -42.5 | -3.365078795603016 | 8 |
| 17.5 | -6.5 | -30.5 | -3.3625912553337383 | 8 |
| 3.5 | -48.5 | -36.5 | -3.3604032223518137 | 8 |
| 21.5 | -70.5 | -56.5 | -3.3517096917008615 | 8 |
| 53.5 | -0.5 | -28.5 | -3.351552596338927 | 8 |
| 69.5 | -44.5 | -0.5 | -3.328113542914116 | 8 |
| -46.5 | -52.5 | -26.5 | -3.3244978184003897 | 16 |
| 1.5 | -30.5 | -36.5 | -3.3096372412928043 | 8 |
| -8.5 | 71.5 | 15.5 | -3.3086814338894026 | 8 |
| -58.5 | -40.5 | -8.5 | -3.3044317236324474 | 8 |

| | | | | |
|---|---|---|---|---|
| 13.5 | 71.5 | 15.5 | -3.303901685115651 | 8 |
| -34.5 | -0.5 | -8.5 | -3.3019921885542183 | 8 |
| -70.5 | -26.5 | -22.5 | -3.301362882435639 | 8 |
| 21.5 | 61.5 | 1.5 | -3.3013282954235086 | 8 |
| -60.5 | -60.5 | -22.5 | -3.3002817330965657 | 8 |
| -26.5 | -56.5 | -34.5 | -3.288615473204448 | 8 |
| 67.5 | -48.5 | -16.5 | -3.28591091965025 | 8 |
